# Characterizing meniscal calcifications with photon counting-based dual-energy computed tomography

**DOI:** 10.1101/2025.02.14.638040

**Authors:** Eeva Nevanranta, Ville-Pauli Karjalainen, Mikael Brix, Iida Hellberg, Aleksandra Turkiewicz, Bijay Shakya, Patrik Önnerfjord, Sampo Ylisiurua, Amanda Sjögren, Khaled Elkhouly, Velocity Hughes, Jon Tjörnstrand, Simo Saarakkala, Martin Englund, Mikko A.J Finnilä

**Author notes:** **Corresponding author:** Dr. Mikko A.J. Finnilä, Research Unit of Health Sciences and Technology, Faculty of Medicine, University of Oulu, Pob 5000, FI-90014 Oulu, Finland.

## Abstract

Meniscal calcifications are associated with meniscal degeneration and osteoarthritis (OA). However, differentiating between calcification types, such as basic calcium phosphate (BCP) and calcium pyrophosphate (CPP), remains challenging *in vivo*. Therefore, assessing new imaging possibilities is crucial in understanding the calcification pathologies and their relationship with OA. This study investigated dual-energy computed tomography combined with a photon counting detector (PCD-DECT) to identify calcifications in human meniscal samples *ex vivo*, using Raman spectroscopy as reference. 82 meniscus samples from 41 donors were imaged using PCD-DECT at 120kVp. Data were collected in two energy bins (20-50keV and 50-120keV) with an isotropic resolution of 37µm. Raman spectroscopy was used to classify the calcifications as BCP or CPP. Among 82 samples, Raman spectroscopy identified 36 samples with only either BCP or CPP calcifications, that were included in the subsequent analysis. Regression models were used to compare the dual-energy index (DEI) and to assess the low energy values for corresponding high energy values between calcification types. The highest difference observed between BCP and CPP in comparison of low energy values for corresponding high energy values was 166HU (95%CI: 73, 259) at high energy value of 500HU and the difference between DEI values was 0.035 (95%CI: 0.011, 0.059), suggesting a potential difference in the measured parameters of calcification types. To conclude, PCD-DECT allowed the measurement of BCP and CPP calcifications *ex vivo*, offering new potential *in vivo* applications in the future, which could help understand calcification processes and evaluate the efficacy of disease-modifying drugs targeting calcification inhibition.

## 1. INTRODUCTION

The menisci of the human knee are crescent-shaped fibrocartilaginous structures that provide essential biomechanical functions such as load distribution, shock absorption, and joint stability by transmitting weight and reducing friction (MCDEVITT and WEBBER, 1990). Menisci are predominantly composed of water rich well organized extracellular matrix, that with the smooth surfaces and wedge-like cross-sections provide crucial functionality (MCDEVITT and WEBBER, 1990). However, the prevalent disease osteoarthritis (OA) commonly affects the menisci, impairing biomechanics, increasing friction, and compromising joint stability (Loeser et al., 2012; Yao et al., 2023). Additionally, meniscal calcifications, specifically basic calcium phosphate (BCP) which includes hydroxyapatite (HA), and calcium pyrophosphate (CPP) are commonly associated with meniscal degeneration and OA (Fuerst et al., 2009a; Hellberg et al., 2023; Sun et al., 2010; Sun and Mauerhan, 2012). These dense calcification deposits may further contribute to the degeneration and destabilization of the meniscus’ biomechanical functions. The cause-and-effect relationship between meniscal calcifications and OA remains unclear due to the current lack of methods for early detection and monitoring of calcification accumulation in the initial stages of the disease (Fuerst et al., 2009a; Yao et al., 2023). Identifying the calcification types *in vivo* could potentially aid in assessing their role in OA development.

Previously, accurate methods to identify the types of intra-articular ectopic calcifications have included Raman spectroscopy, scanning electron microscope, and X-ray diffraction (Hawellek et al., 2018; Katsamenis et al., 2012). However, these methods can only be used *ex vivo* and require either histological sections with exposed calcification or tissue samples, necessitating access to meniscal tissues, which is only possible from end-stage OA patients or cadavers but not *in vivo*. Therefore, clinical studies have explored dual-energy computed tomography (DECT) as a potential modality to identify calcification types *in vivo*; DECT has provided promising results in differentiating BCP calcifications from CPP deposits (Pascart et al., 2020a, 2019). However, the results and accuracy between different studies have varied, and opposing views on the efficacy of DECT in calcification identification have also been proposed (Døssing et al., 2021; Jarraya et al., 2024).

Photon counting detector (PCD) enables the collection of more than two energy bins in a single scan, resulting in the multi-energy computed tomography (MECT) modality (Willemink et al., 2018). This advancement enhances the accuracy of both *ex vivo* and *in vivo* imaging compared to conventional DECT, enabling the better quantification of various materials by resolving material-specific attenuation functions (Becce et al., 2019; Bhayana et al., 2020). CT devices with PCDs are already in clinical use (Greffier et al., 2023a; Holmes et al., 2023) and with the ability to capture multiple energy levels simultaneously, it could also be used to distinguish between different types of calcifications based on their unique attenuation profiles and lower radiation dosage compared to DECT (McCollough et al., 2015). Moreover, PCDs enhance the differentiation capability by providing higher spatial resolution, lower electronic noise, and simultaneous multi-energy imaging without increasing the radiation dose (Greffier et al., 2023a; McCollough et al., 2023, 2015). Recent MECT studies have shown promising results for differentiating calcification types (Becce et al., 2019; Stamp et al., 2019). However, the research has been restricted to limited comparisons of calcifications from different tissue sources, small sample sizes, or spatial resolution.

In this study, we investigate the efficacy of DECT combined with PCD, hereafter referred to as PCD-DECT in identifying meniscal calcification deposits in the posterior horns of human menisci *ex vivo*, using Raman spectroscopy as a reference from our previous study (Shakya et al., 2024). We aim to differentiate between the calcification types with a high-resolution PCD-DECT device using a large sample set consisting only of meniscal tissue.

## 2. METHOD

### 2.1. Tissue samples

This study was approved by the regional ethical review board at Lund University (Dnr 2015/39 and Dnr 2016/865). The same sample set was used in our previous study where the sample selection is explained in more detail (Shakya et al., 2024). Briefly, the human meniscus samples were sourced from the knee tissue biobank MENIX, based in Skåne University Hospital in Lund, Sweden. For this study, a total of 82 meniscus samples were collected. The collection included medial and lateral menisci from a single knee of 21 deceased adult donors without a history of knee OA or rheumatoid arthritis and 20 medial compartment knee OA patients who underwent a total knee replacement (TKR). The presence of medial compartment OA was assessed based on the surgeon’s Outerbridge classification of knee joint cartilage. TKR patients with a medial grade IV and a lateral grade lower than IV were included in the study. All samples were preserved at -80 degrees Celsius within two hours of extraction.

### 2.2. Sample preparation

Upon receiving the meniscus samples from the biobank, they were thawed in phosphate-buffered saline (PBS) and divided into two sections (Figure 1). The whole posterior horn section was used for this study. The posterior horn samples were then fixed in 4% saline-buffered formaldehyde for at least 11 days before undergoing PCD-DECT scanning. Following the scanning, the samples were processed histologically to paraffin blocks. Additionally, in our previous study, 4-µm-thick slices were prepared for Raman measurements to characterize the calcification types. The meniscus samples were divided into BCP and CPP calcification type groups based on the Raman measurements (Shakya et al., 2024).

**Figure 1.**
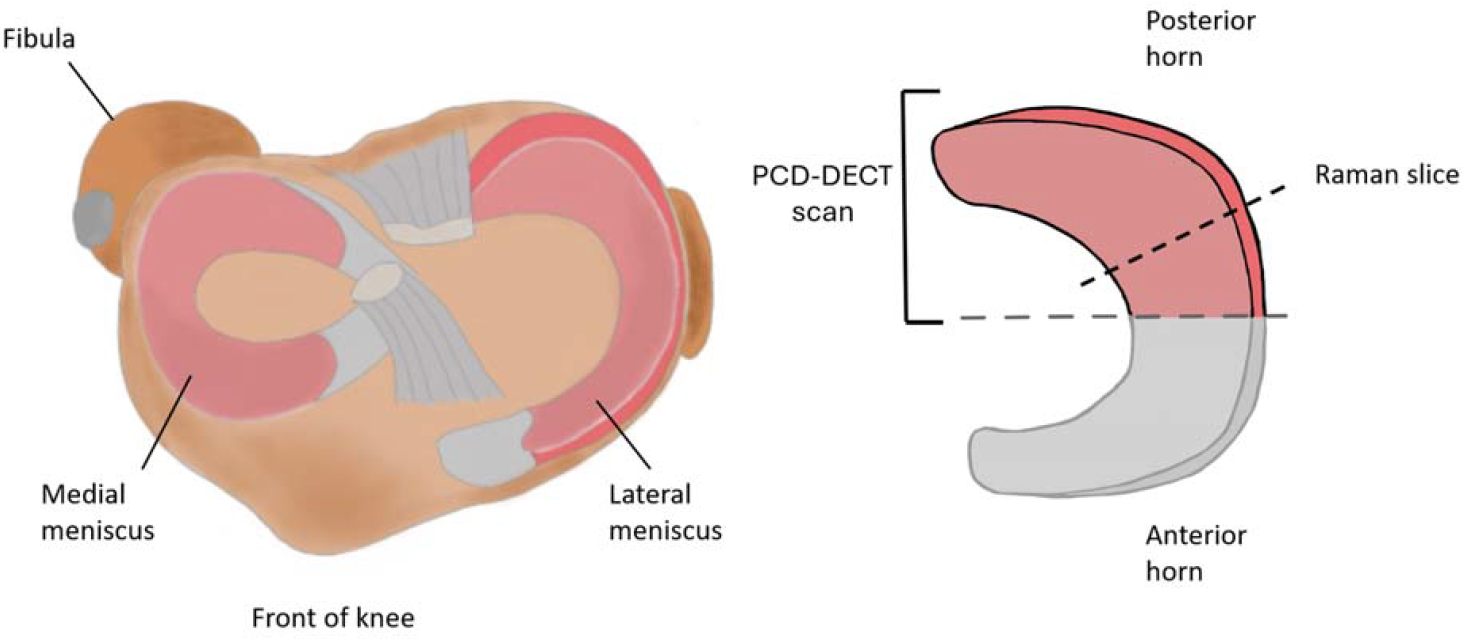
A) An illustration of medial and lateral menisci in the knee joint. Both the medial and lateral posterior horns were included in this study. B) The meniscus samples were cut into two pieces; the piece including the posterior horn was used in this study. The highlighted part of the posterior horn of the meniscus was then imaged with a dual-energy computed tomography (DECT) device coupled with a photon counting detector (PCD). After the imaging, the meniscus was sectioned into smaller segments and 4-µm-thick slices were cut and prepared for Raman spectroscopy.

### 2.3. Photon counting based dual-energy computed tomography imaging

The meniscus posterior horn samples were scanned in the Medical Imaging Teaching and Test Laboratory (MITTLAB) using an experimental cone-beam CT setup, which includes an X-ray tube, a PCD, and a motorized rotator. The PCD (Flite FX15, XCounter AB, Danderyd, Sweden) is a tilted flat-panel detector with a 5.13 × 15.47 cm^2^ active area that uses CdTe for direct conversion. It has two adjustable energy thresholds which were set to 20 keV and 50 keV resulting in a low energy bin of 20 keV-50 keV and a high energy bin of 50 keV-120 keV. The X-ray tube had a 50 µm focal spot and was operated at 120 kVp with a current of 0.2 mA. The sample rotated continuously on a rotation stage (NR360S/M, Thorlabs, Inc., Newton, New Jersey), with the detector and source stationary, at an angular velocity of 0.2038°/s and a frame rate of 3 projections/s, collecting 5300 projections during the 360° scan for a total of 29-minute scan time. The source-detector distance was 86.16 cm, and the sample-detector distance was 50.5 cm, resulting in a final voxel size of 37 µm. Samples were rinsed in PBS and enclosed in parafilm before imaging to prevent drying and movement during the imaging process. Separate calibration scans involving an air scan, and 5.3 cm and 10.6 cm thick PMMA plate scans to calculate signal-to-equivalent thickness correction (STC) were performed (Juntunen et al., 2020). Image reconstruction utilized modified MATLAB code from Juntunen et al. (Juntunen et al., 2021), incorporating both ASTRA and SPOT toolboxes, with iterative penalized least squares (PWLS) with total variation (TV) regularization algorithm. The PWLS with TV reconstruction method was employed, using Barzilai-Borwein iterative step size update with a maximum of 60 iterations, regularization parameter β set to 5, and ε set to 1e-8 to prevent division by zero. Ring artifacts, posing challenges during calcification analysis, were reduced using an algorithm by Münch et al. (Münch et al., 2009) with σ=4 width, followed by inverse FFT transformation for image reconstruction.

### 2.4. Calcification characterization analysis

For the analysis of calcifications, Dragonfly Software (Version 2022.2, ORS Object Research System) was utilized to gather and evaluate the data. First, we co-registered the Raman slice with the PCD-DECT data. After this, as the co-registered PCD-DECT section showed consistent average dual-energy index (DEI) values and similar histograms with the calcifications of the whole sample, we used the Raman result as a reference for the whole PCD-DECT sample. Then, we used global thresholding to remove soft tissue and segment the calcifications, and after analyzed individual calcifications separately. Lastly, for each individual calcification the low (20keV-50keV) and high energy (50keV-120keV) intensities were collected, and the (DEI) (Graser et al., 2008) values were calculated for each individual calcification.

### 2.5. Statistical analysis

Stata (Version 18.5) was used for statistical analysis of low and high energy variables. We used low energy as an outcome. The independent variables were high energy, group (CPP vs BCP), interaction between group, and high energy, high energy squared, interaction between high energy squared and group. The interaction terms are included to allow the associations to differ by group. We have included random intercept for persons and for meniscus (menisci nested within persons). Due to some heteroscedasticity in the data, we used robust standard errors. We evaluated the model fit using residuals diagnostic plots and no model violations were detected. We removed 0.7% of data points with extreme values (high energy <-200 or >1000, low energy >2000) as they were judged to be due to image artifacts. For analysis, we used the command mixed in Stata. To illustrate results, we used Stata commands: margins and margins plot to extract between group differences in low energy values for a range of high energy values. Estimates are presented with 95% confidence intervals. For DEI variable, we did a group-wise comparison of estimated means between BCP and CPP groups using Linear Mixed Models in SPSS (IBM SPSS Statistics v. 29.0, NY, USA). For the comparison of the mean DEI values between BCP and CPP groups, DEI was set as a dependent variable, and we used random intercept to account for calcifications from the same meniscus and a second random effect to account for multiple menisci from the same knee. The descriptive data in the figures are presented as means with 95% CIs. In addition, the number of analyzed individual calcifications in each sample are seen in Supplementary Material Figure S1.

## 3. RESULTS

### 3.1. Description of study subjects

The average age of the individuals with OA and deceased donors was 70 years, and half of the study subjects were female (Table 1).

**Table 1.**
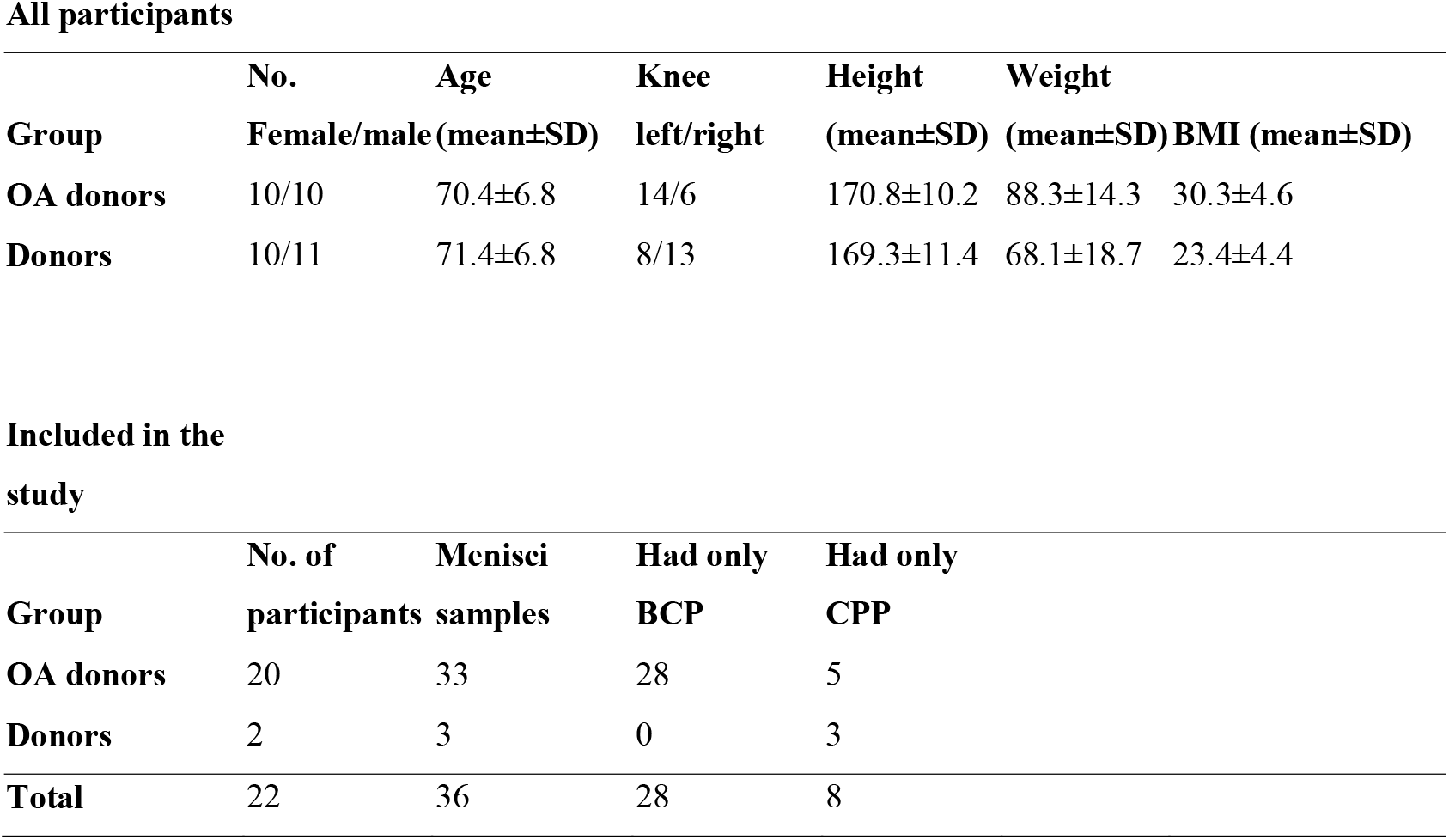
Descriptive statistics of study participants.

All participants
| Group | No.<br>Female/male | Age<br>(mean±SD) | Knee<br>left/right | Height<br>(mean±SD) | Weight<br>(mean±SD) | BMI (mean±SD) |
| --- | --- | --- | --- | --- | --- | --- |
| OA donors | 10/10 | 70.4±6.8 | 14/6 | 170.8±10.2 | 88.3±14.3 | 30.3±4.6 |
| Donors | 10/11 | 71.4±6.8 | 8/13 | 169.3±11.4 | 68.1±18.7 | 23.4±4.4 |

Table 1. Descriptive statistics of study participants.
| Group | No. of<br>participants | Menisci<br>samples | Had only<br>BCP | Had only<br>CPP |
| --- | --- | --- | --- | --- |
| OA donors | 20 | 33 | 28 | 5 |
| Donors | 2 | 3 | 0 | 3 |
| **Total** | 22 | 36 | 28 | 8 |

### 3.2. Calcification characterization with PCD-DECT

In our previous study, from the 82 menisci, we identified 8 samples with CPP type calcifications and 28 samples with BCP type calcifications with Raman spectroscopy, and these 36 samples, divided into CPP and BCP groups, were used in PCD-DECT analyses (Figure 2).

**Figure 2.**
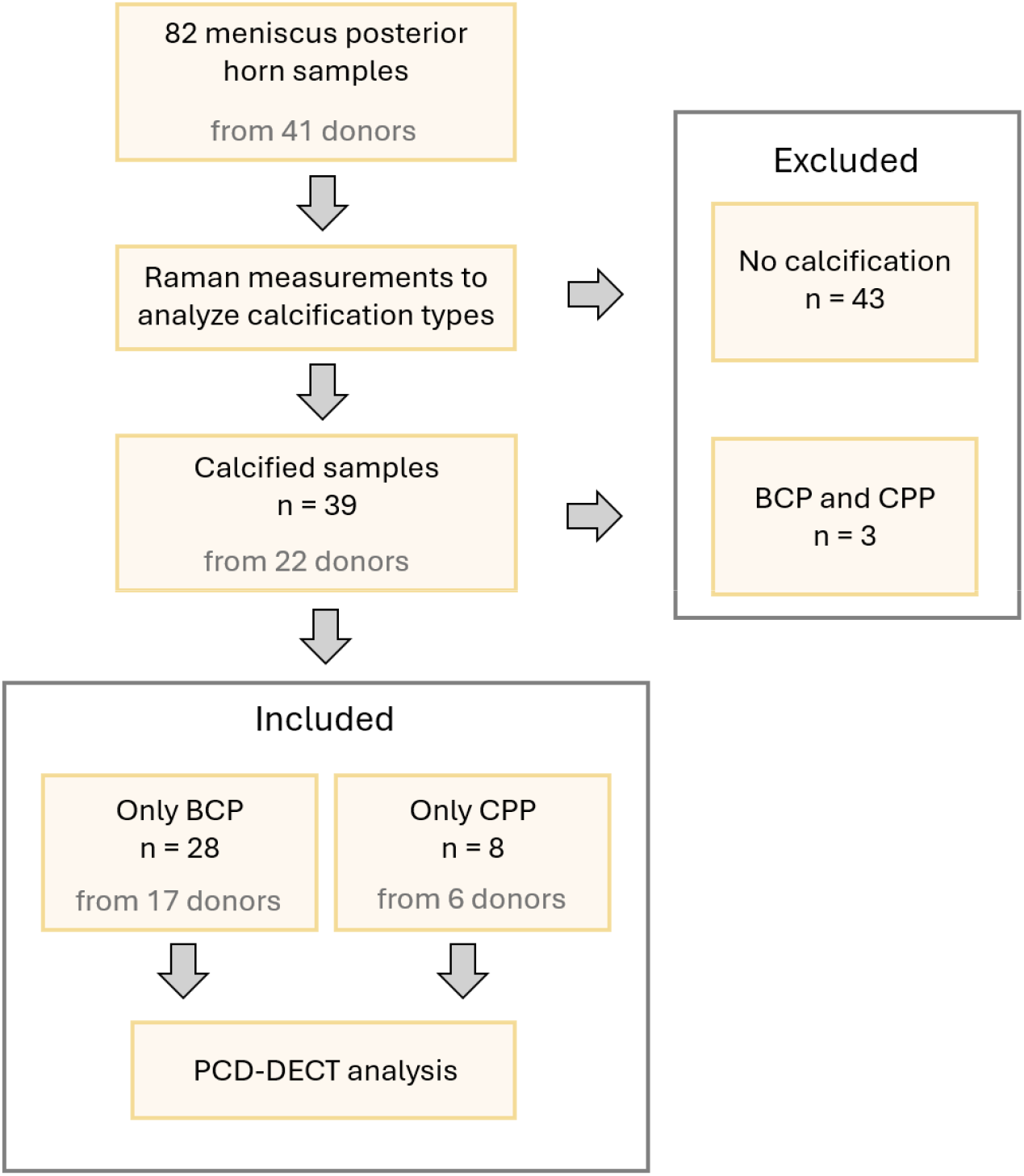
Flow chart presenting excluded and included menisci in the main analysis. A total of 28 BCP samples were collected from 17 different patients, with 16 samples from the medial side and 12 from the lateral side. CPP samples were collected from 6 different patients, with 5 samples from the medial side and 3 from the lateral side. One OA donor inhibited BCP in one meniscus and CPP in the other. In addition, we identified 3 meniscal samples that contained both types of calcifications that were left out from further PCD-DECT analyses.

Subsequentially, all individual calcifications were analyzed in either BCP or CPP groups. We identified a total of 9898 calcifications from the 36 meniscus samples that were included in the analysis. Representative images of samples without calcification, with identified CPP, and with identified BCP calcifications are seen in Figure 3.

**Figure 3.**
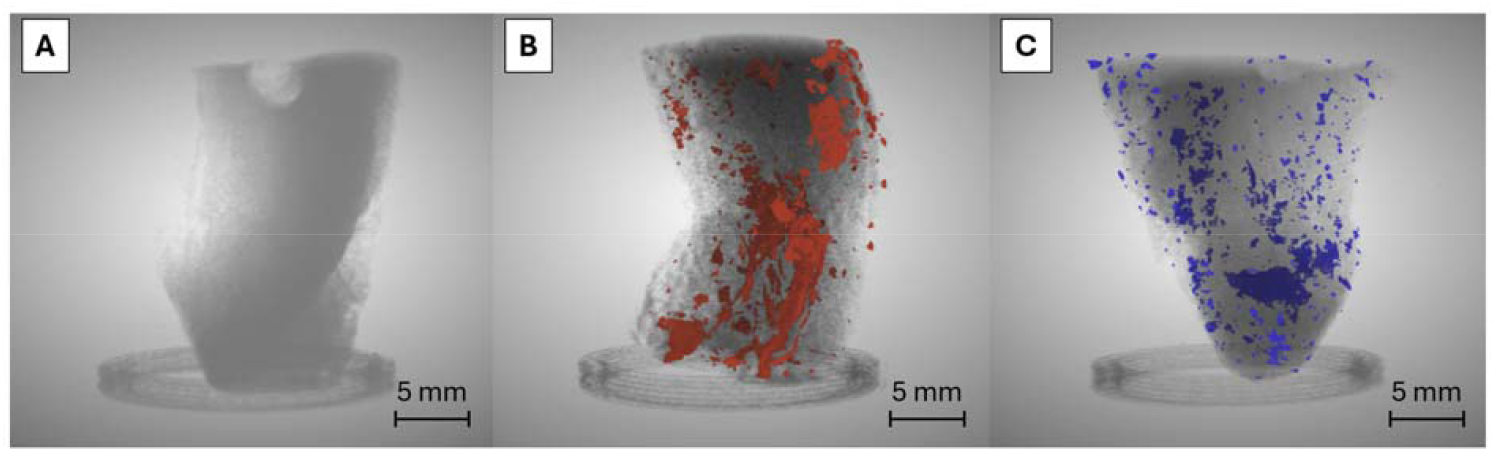
Representative PCD-DECT 3D volumes from three different samples. The calcification types were characterized with Raman spectroscopy A) An example of intact meniscus without any calcification. B) Calcium Pyrophosphate (CPP) calcifications are highlighted in red. C) Basic Calcium Phosphate (BCP) calcifications are highlighted in blue.

The curves in Figure 4 represent the attenuation differences between low and high energy values for CPP and BCP calcifications. The results indicate a difference between CPP and BCP across various high energy values, with CPP showing lower low energy values for a given high energy. The differences are relatively small at lower high energy values and the confidence intervals overlap. However, as the high energy values increase, the difference becomes more pronounced, with the largest difference observed at a high energy value of 500 HU, where the difference is 166.11 HU (95% CI: 73.39, 258.84) HU. This shows the highest difference between the two crystal types. The differences remain evident through higher high energy values, although the magnitude of the difference decreases slightly as the high energy values increase beyond 600 HU. The smallest difference occurs at -100 HU, where the difference is 33.81 HU (95% CI: -40.38, 107.99) HU. Overall, for most points, the difference is around ∼ 50 HU to ∼ 250 HU while the greatest difference between CPP and BCP was observed to be around a high energy value of 500 HU. The detailed differences in specific high energy values are provided in the Supplementary Figure S2.

**Figure 4.**
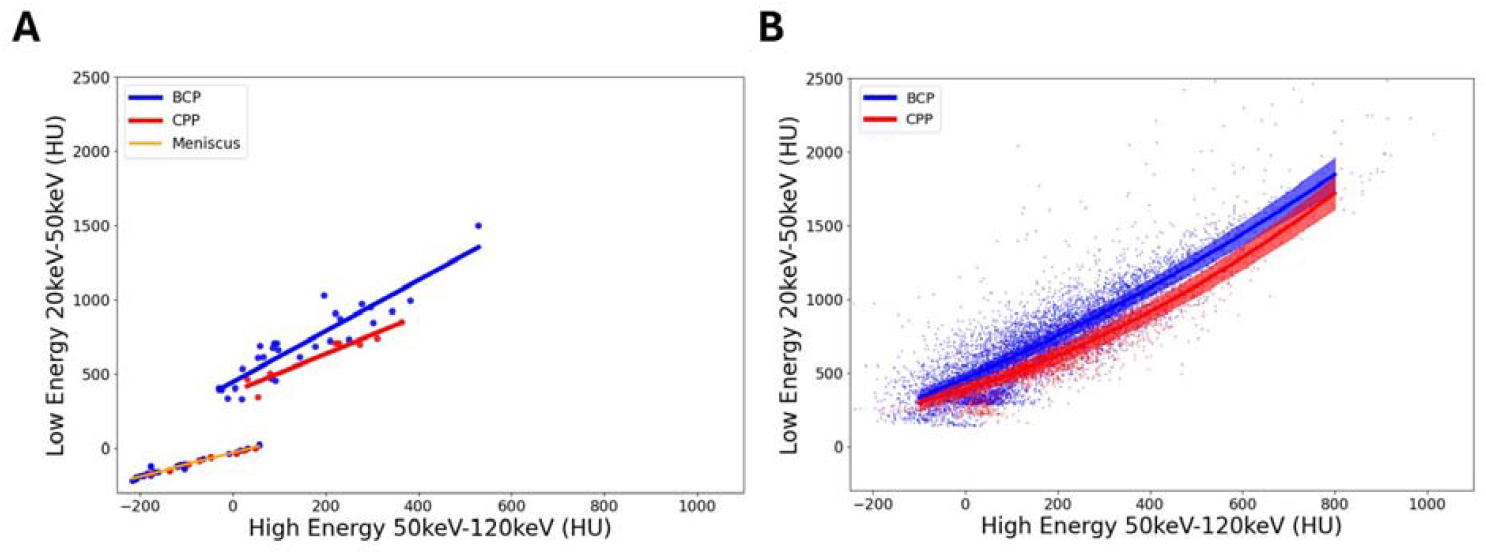
The mean low energy/high energy values per each meniscus with a fitted line for BCP (blue) and CPP (red) calcification groups were calculated from the Hounsfield Unit (HU) value of each sample’s calcifications. In addition, we show the linear association of low and high energy values of meniscal soft tissue from all 36 analyzed samples in yellow line with samples colored blue (BCP) and (CPP) to correspond to the calcification type found in that specific sample. B) All individual calcifications from all 36 samples, presented with high energy values as the reference and corresponding low energy values, including regression fits for both groups to illustrate average distributions, along with 95% confidence intervals.

In the statistical comparison of DEI values between individual BCP and CPP calcifications, we observed higher DEI values of 0.189 (95% CI 0.177, 0.200) in the BCP group compared to 0.154 (95% CI 0.133, 0.175) in CPP group (Table 2).

**Table 2.** The mean dual-energy index (DEI) within all identified BCP and CPP calcifications, and the estimated differences between the BCP and CPP groups. Additionally, we show the soft tissue DEI.

| **Group** | **No. of samples** | **No. of identified calcifications** | **Dual Energy Index (DEI) (95% CI)** | **Estimated difference between BCP and CPP (95% CI)** |
| --- | --- | --- | --- | --- |
| **BCP** | 28 | 7152 | 0.189 (0.177, 0.200) |  |
| **CPP** | 8 | 2746 | 0.154 (0.133, 0.175) |  |
|  |  |  |  | 0.035 (0.011, 0.059) |
| **Soft tissue** | 36 |  | 0.037 (0.023, 0.050) |  |

## 4. DISCUSSION

In this study, we employed an experimental PCD-DECT to detect and characterize the BCP and CPP calcification deposits in the posterior horn of the human meniscus. We observed variation in low energy values for corresponding high energy values between BCP and CPP calcifications, with the regression curves exhibiting slightly different shapes and BCP showing consistently higher low energy values than CPP. Similarly, we observed higher DEI values in the individual BCP calcifications when compared to CPP calcifications. To our knowledge, this is the first study to examine differences in BCP and CPP type calcifications in the meniscus using PCD-DECT from a large sample set consisting of only meniscal samples.

From our calcification analyses, we observed higher DEI values in meniscal BCP calcifications compared to CPP calcifications. Similar results were found in previous studies using DECT where HA in bone was reported to have higher DEI than CPP in meniscus (Pascart et al., 2019), as well as higher DEI in HA compared to CPP in synthetic phantoms of equal concentrations (Pascart et al., 2020a). However, another study reported that DECT has limited potential in differentiating BCP and CPP calcifications (Jarraya et al., 2024). Our *ex vivo* measurements of BCP and CPP crystals suggested a clear distinction at the group level when using the PCD-DECT device. Nevertheless, at the level of individual calcifications, we observed overlap in DEI values, indicating that precise differentiation between individual BCP and CPP calcifications remains challenging with current imaging methods. A possible explanation for varying DEI values between our and previous studies could be the different concentrations and sizes of measured calcifications in the tissue. DEI values have been shown to increase with increasing calcification concentrations in the synthetic calcification phantoms (Budzik et al., 2021; Pascart et al., 2020a). In a biological environment, having only a single concentration of material of interest is unlikely, as even BCP is a broader term encompassing a group of calcium phosphates, including HA (Fuerst et al., 2009b; Stack and McCarthy, 2016), and other calcium containing crystals like calcite have been reported to be commonly found in the knee joints of OA patients (Niessink et al., 2024). Furthermore, in a recent DECT study, it is suggested that the calcium concentration of the calcification deposits affects the DECT parameters more than their biochemical composition (Jarraya et al., 2024). However, we reported in our previous study that this density dependence is related to the STC technique required for the PCD panel when using PCD-DECT device (Juntunen et al., 2020). The STC approach assumes that the beam hardening and scattering properties are identical to those of the calibration material (PMMA), leading to overcorrection of scattering. This overcorrection can introduce a thickness or density dependence in the DEI value estimation, which was also evident in our study. This is further relevant when measuring calcifications from multiple patients or samples affected by different disease pathologies. Therefore, we also examined the low energy values for corresponding high energy values across the calcification types. We observed that the BCP group generally exhibited higher low energy values than the CPP group for a given high energy value. As density alone should not affect DEI values the variability comes mostly from beam hardening and resolution. Additionally, a higher count of CPP calcified samples could provide a more accurate estimation of the low and high energy regression and DEI values and is critical for any kind of generalization of the results. Therefore, in future studies focusing on the characterization of calcifications, reporting both DEI values and the low energy values for corresponding high energy values for each calcification type is highly recommended.

In our study, in the PCD-DECT scans we employed a lower energy level (20keV - 50 keV and 50keV - 120 keV) and a higher resolution (37 µm voxel size) than in typical clinical DECT settings (*i.e*. 80 and 140 kVp) (Bhayana et al., 2020; McCollough et al., 2015; Rajiah et al., 2020). A previous MECT study identified more attenuation differences for BCP and CPP calcifications using similar low energy ranges (Becce et al., 2019). Another study suggested that the identification of calcifications benefits from having more data points within the region of interest, which can be achieved with higher resolution (Døssing et al., 2021). Therefore, using lower energies and higher resolution than those typically employed in clinical DECT and MECT devices would enhance the detection of calcifications (Becce et al., 2019; Stamp et al., 2019). However, in clinical settings, this approach is not feasible due to significant beam hardening from overlapping anatomical structures. Moreover, higher resolution can also minimize the measurement error from partial volume effect (Soret et al., 2007), which is a significant factor complicating the identification of calcifications; it arises when the size of the calcifications is much smaller (<20µm) (Yavorskyy et al., 2008) compared to the imaging resolution in clinical devices (∼0.5mm^3^) (Greffier et al., 2023a). A recent CT study showed that measuring a material of interest in different background mediums caused large errors in the elemental mass assessment; these errors are dependent on the varying attenuation coefficient of the different background mediums (Salyapongse and Szczykutowicz, 2024). This is relevant for our study, as the intensity values of calcium phosphates i.e. the calcification lies between soft tissue and bone, measuring the same concentration of calcification in soft tissue would have decreased DEI value when compared to bone as a background medium, which would largely affect the measurement result. Thus, in our study, we only used meniscal tissue as a background medium for both CPP and BCP calcifications. In future studies, when differentiating constituents like BCP and CPP calcifications, we suggest that the measurements are done only with a single tissue type as background medium because it decreases the measurement variability and leads to more accurate material identification.

Raman spectroscopy, which was conducted in our previous study for this sample set, was used as a reference method to divide the samples into BCP and CPP groups (Shakya et al., 2024). However, Raman spectroscopy can only analyze a single 4-µm-thick section at a time requiring another method to cover larger volumes of interest. In our study, after co-registering the PCD-DECT and Raman samples and analyzing the same area in each sample using both methodologies, consistent results were shown when compared to the PCD-DECT results of whole sample measurements. This further validated the use of whole samples during PCD- DECT analyses. Compared to Raman spectroscopy and conventional histology as an identification method, PCD-DECT is advantageous in providing 3D data and therefore covering larger volumes of interest non-destructively. Furthermore, it enables *in vivo* imaging of calcifications in soft tissues (Pascart et al., 2020b). In our study, PCD-DECT identified a few samples with minute calcifications that could not be located in histological sections or Raman spectroscopy, suggesting that high-resolution PCD-DECT is a more sensitive method compared to methods requiring conventional histological sectioning. Despite this advantage, most DECT devices currently operate with a resolution of 0.2-0.6mm, leading to significant errors due to the partial volume effect, as the calcification particles are generally much smaller than the voxel size (Greffier et al., 2023b). The Naeotom Alpha, currently the only commercially available PCD-DECT device, offers the best resolution with 0.11 mm and a slice thickness of 0.2 mm (Vecsey-Nagy et al., 2024). However, even this resolution remains much larger than the typical size of calcification particles, may limit its ability to resolve these structures accurately. Therefore, PCD-based technologies still have potential for improvement through advancements in detector technology, which could further enhance material differentiation *in vivo*. To observe smaller particles, the resolution should be improved with even smaller pixel sizes, and enhanced calibration techniques could contribute to more precise tissue quantification and material characterization. Additionally, the addition of more energy channels could provide more accurate information from attenuation changes, leading to better differentiation of materials.

It is important to bear in mind the limitations of our study. First, the HU values in this study are not directly comparable to clinical DECT values due to the lower energy levels used than in typical clinical settings. However, a similar problem also persists between different MECT and DECT devices since HU values are not interchangeable with other devices, and the relative differences between the BCP and CPP types are still valid for comparison. Additionally, the sample grouping into BCP and CPP categories was based on thin sections studied with Raman spectroscopy, that account only a small portion of the entire posterior horn of the meniscus. Therefore, it is possible that a small number of other calcification types are in the sample that are not seen in Raman sections. However, the PCD-DECT analysis showed similar average DEI values in both the whole sample and when compared to the co-registered Raman section, suggesting that small traces of other calcification types are averaged out in the final analysis.

In conclusion, the PCD-DECT analysis indicates differences between BCP and CPP calcifications in the measured DEI and in low energy values for corresponding high energy values in posterior horns of human menisci *ex vivo*. PCD may offer new potential in future *in vivo* applications that could help understand the calcification processes in the knee joint and help evaluate how disease-modifying drugs could inhibit calcifications. While the differentiation ability of PCD-DECT can be further improved in the future with advancing technologies, the efficacy of the device can also decrease if aspects like varying densities, different calcification tissue sources, and resolution are not taken into careful consideration during measurements and analysis. Failing to account for previously mentioned aspects is the most likely reason for converging or overlapping results when differentiating the calcification types. We conclude that in future studies, these aspects should be considered carefully.

## ACKNOWLEDGEMENTS

We would like to thank the MENIX clinical staff at Trelleborg Hospital, the Tissue Donor Bank at Skåne University Hospital, and the Department of Forensic Medicine in Lund for their collaboration that enabled sample collection. We would also like to acknowledge Piia Mäkelä, Laboratory Technician, for the preparatory work with the histological samples.

## AUTHOR CONTRIBUTIONS

Conception and design: EN, VPK, IH, AT, SS, ME, MF Analysis and interpretation of the data: EN, VPK, MB, AT, MF Drafting of the article: EN, VPK

Critical revision of the article for important intellectual content: All co-authors Final approval of the article: All co-authors

Provision of study materials or patients: ME, VH, PÖ, JT, VPK, KE, AS, SS Collection and assembly of data: EN, VPK, MB, SY

