## Supplementary figures and images for "Characterizing meniscal calcifications with photon counting-based dual-energy computed tomography"

### Supplementary Table 2

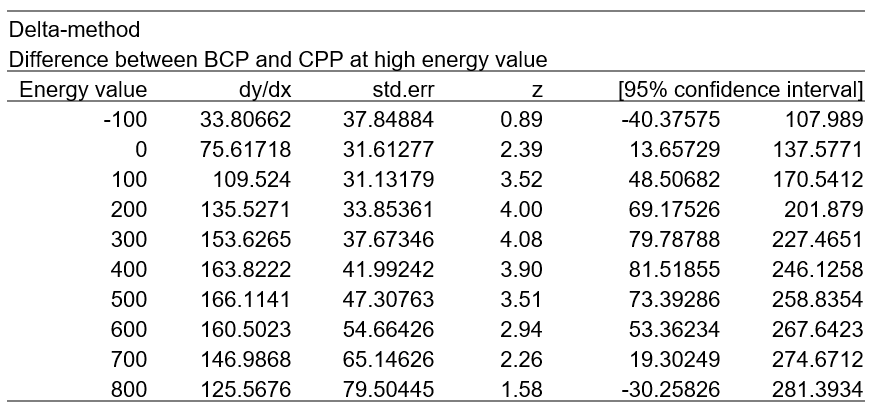

**Figure S2**. The difference between BCP and CPP at high energy value.
