## Supplementary Table 1 for "Characterizing meniscal calcifications with photon counting-based dual-energy computed tomography"

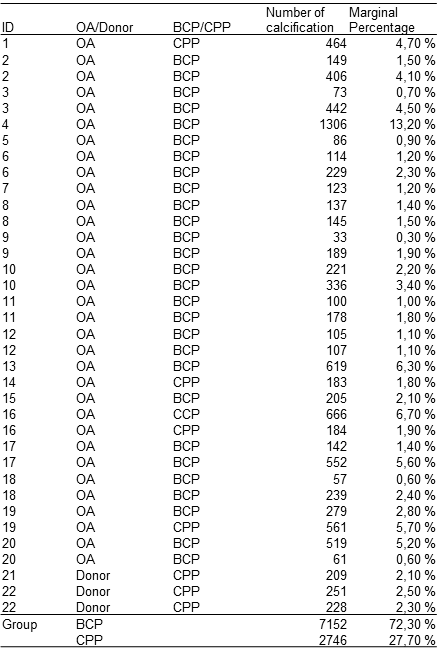

**Figure S1.** The number of analyzed calcifications used in the study, along with their corresponding OA group and calcification type.
